# Evolutionary conservation of A/T-ending codons reflects co-regulation of expression and complex formation

**DOI:** 10.1101/2022.01.17.475622

**Authors:** Hannah Benisty, Xavier Hernandez-Alias, Marc Weber, Miquel Anglada-Girotto, Federica Mantica, Leandro Radusky, Gökçe Senger, Donate Weghorn, Manuel Irimia, Martin H. Schaefer, Luis Serrano

**Affiliations:** Centre for Genomic Regulation (CRG), The Barcelona Institute of Science and Technology, Dr. Aiguader 88, Barcelona 08003, Spain; Department of Experimental Oncology, IEO, European Institute of Oncology IRCCS, Via Adamello 16, Milan 20139, Italy; Universitat Pompeu Fabra (UPF), Barcelona, Spain; ICREA, Pg. Lluis Companys 23, Barcelona 08010, Spain

**Author notes:** Correspondence (L.S.), (H.B.).

**Keywords:** Codon bias, Conservation, Mammals, tRNA

## Abstract

**Background:** In a wide variety of organisms, synonymous codons are used with different frequencies, a phenomenon known as codon bias that plays an important role in determining expression levels. However, the importance of codon bias to facilitate the simultaneous turnover of thousands of protein-coding transcripts to bring about phenotypic changes in cellular programs such as development, has not yet been investigated in detail.

**Results:** Here, we discover that genes with A/T-ending codon preferences are expressed coordinately and display a high codon conservation in mammals. This feature is not observed in genes enriched in G/C-ending codons. A paradigmatic case of this phenomenon is KRAS, from the RAS family, an A/T-rich gene with a high codon conservation (95%) in comparison to HRAS (76%). Also, we find that genes with similar codon composition are more likely to be part of the same protein complex, and that genes with A/T-ending codons are more prone to form protein complexes than those rich in G/C. The codon preferences of genes with A/T-ending codons are conserved among vertebrates. We propose that codon conservation, a feature of expression-coordinated transcripts, is linked to the high expression variation and coordination of tRNA isoacceptors reading A/T-ending codons.

**Conclusions:** Our data indicate that cells exploit A/T-ending codons to generate coordinated, fine-tuned changes of protein-coding transcripts. We suggest that this orchestration contributes to tissue-specific and ontogenetic-specific expression, which can facilitate, for instance, timely protein complex formation.

## Background

In all species of life, the same amino acid can be encoded by up to six different synonymous codons. The preference for certain synonymous codons over others is referred to as codon bias. This phenomenon is widely observed between species and between genes in a genome, and the reasons for the distinct patterns of codon usage have been debated for decades [1, 2]. Explanations involve either mutational or selective causes, which are not mutually exclusive [3, 4]. Synonymous variations are often assumed to be selectively neutral because they do not alter the sequence of the encoded protein [5]. However, the most efficiently translated codons are those matching the tRNAs with the most abundant copies in the cell [6]. This observation has traditionally supported the idea that codon usage could have a fitness effect and, hence, should be under selection rather than only being the result of mutational biases [3, 7]. Even though selection remains highly debated, especially for multicellular species [8, 9], evidence supports the encoding of additional layers of information by an ensemble of codon-mediated mechanisms. For instance, selection may arise from local requirements imposed by mRNA processing. In this sense, alternative splicing regulation motifs have been shown to be under selective pressure at synonymous codons [10, 11]. In addition, codon usage acts on mRNA stability [12], mRNA localization [13], mRNA translation speed and accuracy [14], or protein folding [15, 16]. Also, a growing body of evidence suggests that synonymous mutations in human diseases can have effects that drive selection through their impacts on gene expression [17–19].

Protein-coding transcripts that play a role in cell division are coordinated in their expression and have a codon usage adapted to the tRNA abundances of proliferating cells [20–22]. The realization that specific codon usage is required for dynamic and coordinated changes ensuring the suitable production of certain gene products has led to the expectation that some codon signatures will be conserved along evolution. Indeed, regions of mammalian genes have been described to be under synonymous codon selection [23, 24]. In this sense, we explored here the conservation of codon usage across 57 mammals and its relation with protein-coding transcript regulation. When we looked at codon conservation in HRAS and KRAS, which have an antagonistic codon usage [25], for conserved amino acids, we found that codons of KRAS rarely change in mammals, whereas this is not the case for HRAS. This observation raised the question of whether or not this was a general phenomenon that applied to other genes and, if so, what could be the underlying reasons. We found that the conservation of codons between mammals was related to within-genome variation in codon bias, *i.e.* genes with similar codon usage tend to display a comparable level of codon conservation. We also observed genes with high and low codon conservation, and the main difference between them was that coding sequences with conserved codons were enriched in A/T-ending codons. Then, we found that genes enriched in A/T-ending codons had coordinated changes in mRNA levels across tissues and developmental stages and tended to belong to the same protein complexes. We also investigated whether the highest coordination of transcripts enriched in A/T-ending codons could be related to a specific regulation of tRNA isoacceptors involved in the translation of A/T-ending codons. Additionally, we explored whether the polarized codon usage patterns were also observed in other species of vertebrates and invertebrates.

Altogether, our work highlights a systematic synonymous codon conservation across mammals for protein-coding transcripts that present coordinated changes in expression. Therefore, we speculate that the specific regulation and selection of A/T-ending codons is a mechanism that allows regulating a group of transcripts that need to be expressed coordinately for specific cellular processes, for instance, tissue-specific expression, cell-cycle dependent gene expression, or protein complexes formation.

## Results

### The codons of RAS genes are differently conserved

The three RAS gene paralogs are a paradigm for codon usage bias, as they code for very similar proteins (85% amino acid identity) by using very different codons [25]. We have previously shown that the codon bias observed in RAS genes leads to their regulated differential expression in specific cellular contexts [21]. The synonymous codon usage bias of the KRAS gene itself is associated with enhanced translation efficiency during cell proliferation [21].

We looked at the synonymous codon conservation (see Methods section) of the RAS genes in 57 mammals. The codon usage was highly conserved for KRAS but not for HRAS, although conservation at the amino acid level was similar. For instance, when comparing the codons used in mice and humans, we found a higher percentage of synonymous changes in HRAS than in KRAS, and NRAS showed intermediate values (Figure 1A). This observation points towards a possible selection for the maintenance of the codons in KRAS. This led us to ask if this feature was common in other gene families and to find the characteristics of genes with conserved versus non-conserved codon usage. To tackle this question, we first selected all human genes with more than 90% of average amino acid identity among eutherian mammals (see Supplementary Table 1 for amino acid identities). Then we aligned the codon sequences of these human genes to the corresponding sequences in all the analyzed species of mammals (Supplementary Table 2). Finally, we determined the average codon conservation for those positions where the amino acid was the same in all selected species. The distribution of the synonymous codon conservation varied from 0.62 to 0.98, with KRAS among the genes with highest codon conservation and HRAS among the least conserved genes (Figure 1B, Supplementary Table 3). Our data indicated that synonymous codons of RAS genes and other mammalian genes were differently conserved across mammals.

**Figure 1:**
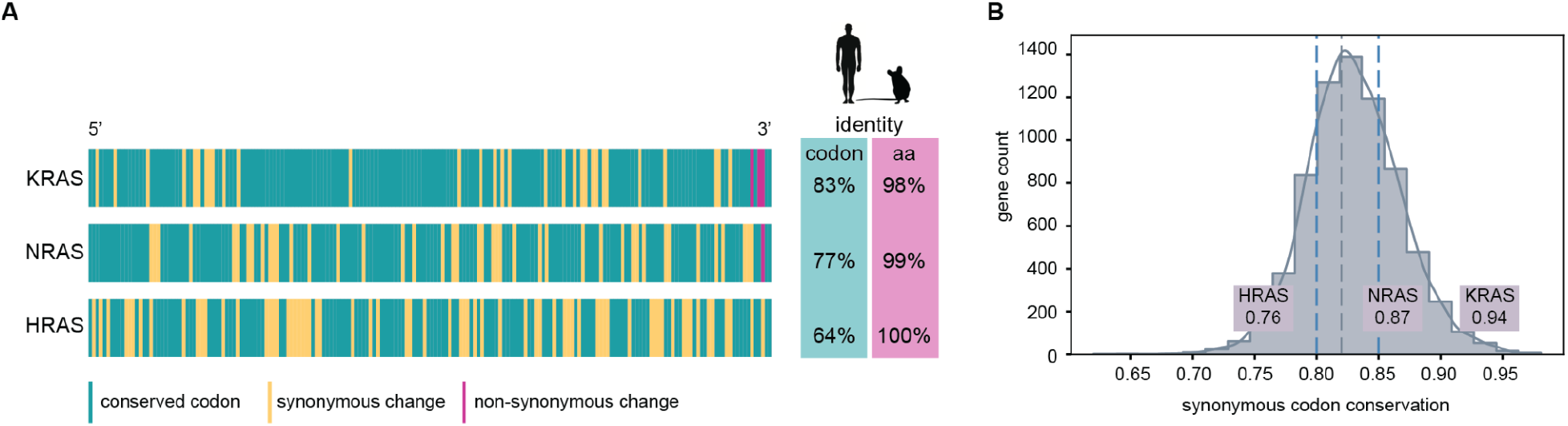
Synonymous codon conservation of coding sequences among mammals. A. Comparison of the codon usage of the RAS coding sequences between human and mouse. Percentages represent the amount of codons conserved (left) and residues conserved (right) between mouse and human when aligning their coding sequences. B. Distribution of the average synonymous codon conservation at position with conserved residues among mammals for all human coding sequences with a high amino acid conservation (>90%). The gray dashed line corresponds to the median, and the blue dashed lines to the 25th and 75th percentile, respectively. The average codon conservation values for RAS coding sequences are indicated.

### Codon usage and conservation among human protein coding sequences

We have previously shown that KRAS is enriched in A/T-ending codons and HRAS in G/C-ending codons [21]. Thus, we wondered if genes with conserved codons have a similar bias for A/T-ending codons as KRAS. To visualize how the codon usage of the studied genes correlates with their codon conservation, we applied principal component analysis (PCA) to the relative codon usage frequencies of all individual genes. We then colored each gene according to the average value of codon conservation and this revealed two distinct poles at the extremes of the first principal component of the codon usage PCA (PC1) (Figure 2A). At one end (negative values of PC1), a strong enrichment of genes with codons conserved among mammals was observed, and at the other end (positive values of PC1), the majority of genes with low codon conservation were gathered. When separating genes in five bins of PC1, an anticorrelation between codon conservation and PC1 values (spearman correlation −0.42) was observed. Genes with the most negative PC1 had a significantly higher (Wilcoxon signed-rank test, p≈0 within numerical error) codon conservation than those in the positive PC1 (Figure 2B). Moreover, genes in the positive pole (with low codon conservation) were enriched in G/C-ending codons, whereas those in the negative pole (with high codon conservation) were enriched in A/T-ending codons (Figure 2C; see GC3 content for all human genes in Supplementary Table 4).

**Figure 2:**
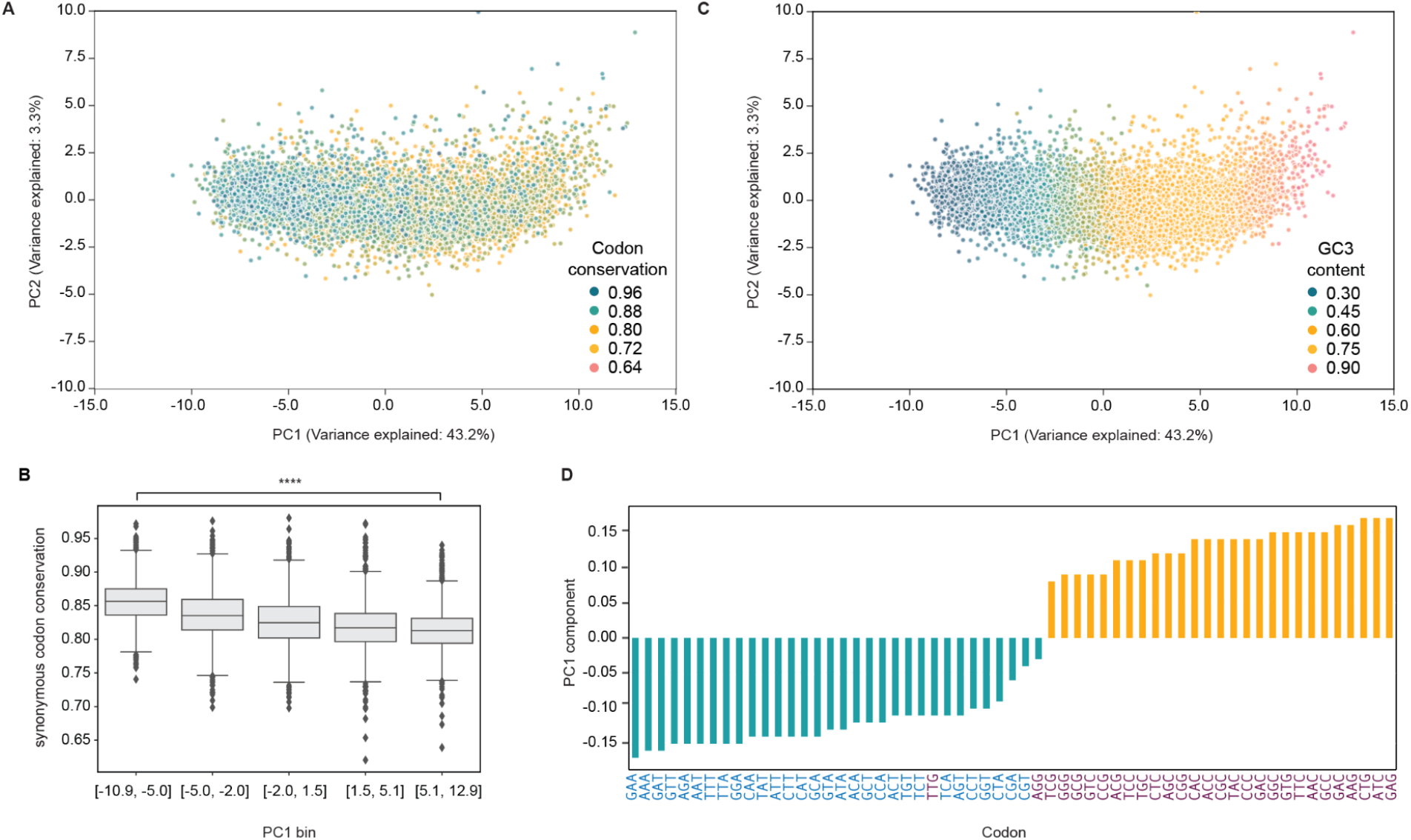
A. PCA projection of the human codon usage. The location of each gene is determined by its codon usage. Distribution of conservation along the main codon usage axis reveals a polarization of codon conservation: the genes with the most conserved (negative PC1) and least conserved codon usage (positive PC1). B. Boxplot of the codon conservation of genes of the different PC1 separated in bins. The codon conservation of genes in the negative PC1 is significantly higher than that of the genes in the positive pole (Wilcoxon signed-rank test, p≈0 within numerical error). C. PCA projection of the human codon usage. The location of each gene is determined by its codon usage. Distribution of average GC3 content among mammals along the main codon usage axis. D. Codons are ordered according to their first component coefficient (PC1). Note that the scale of the values is arbitrary, as only the relative values are important (direction of the vector in the multidimensional space). Negative values indicate a negative PC1 and vice versa. A/T-ending codons are highlighted in blue and G/C-ending codons in purple.

Since the main axis of codon usage seemed to be strongly related to codon usage conservation, we defined two groups of genes from the opposite poles of PC1 for the following analyses. For the sake of readability, we called the group of 1,000 genes with the lowest PC1 coordinates (PC1 < −6.0 and an average of synonymous codon conservation of 0.85) the “negative pole” and the group of 1,000 genes with the highest PC1 coordinates (PC1 > 6.2 and an average of synonymous codon conservation of 0.81) the “positive pole” (Supplementary Table 5).

A gene ontology enrichment analysis revealed that the negative pole included genes related to metabolism and RNA and protein modification, whereas the positive pole consisted of genes involved in the regulation of differentiation and development (Supplementary Figure 1A, Supplementary Table 6). To visualize the importance of each codon across the PC1, we ordered the codons according to their coefficient of the first eigenvector (direction of the first component). We observed that A/T-ending codons were mainly present in the negative pole and G/C-ending codons in the positive pole (Figure 2D, Supplementary Table 7). Thus, two main groups of genes with opposite third-codon position nucleotide content were found, one of which was enriched in A/T-ending codons and was characterized by high codon conservation across mammals.

Gene expression patterns are correlated and stable between species [26]. Therefore, we would expect that this observation holds specially for the genes with a conserved synonymous codon usage. Previous comparative studies have classified genes as evolutionarily stable or variable based on the correlation of their gene expression across species [27]. Harnessing this classification, we found that, in the negative pole, the relative amount of stable genes was two-fold higher than the amount of variable genes while no differences were observed in the positive pole (Chi-squared test, p <0.0001) (Supplementary Figure 1B).

We then explored whether the observed conservation could be related to the importance of the gene function. We used genome-scale CRISPR knockout screens from DepMap [28], where a score of 0 corresponds to a gene not essential for cellular proliferation and a negative score to essential genes, with −1 being the median of all essential genes. We found a significantly larger amount of genes that resulted in essentiality scores in the negative pole than in the positive (Wilcoxon signed-rank test, p= 2.9e-10) (Supplementary Figure 1C).

Briefly, genes rich in A/T-ending codons had a higher synonymous codon conservation than those with G/C-ending codons. The synonymous conservation could be related to the necessity for a fine-tuned regulation of certain genes, since, for instance, the expression of genes in the negative pole was correlated with the expression of the orthologs in other mammals and reflected by their essentiality in cell proliferation.

### Coordination of cellular pools of mRNAs

Given that genes commonly function together, conserved codon usage may reflect concerted expression changes of distinct sets of genes. To identify such expression shifts, we explored the expression from the two groups of studied genes. Transcript abundance is codon-usage dependent [12, 21, 29–31], since ribosome translocation is faster through optimal codons when their cognate tRNAs are abundant and less prone to be degraded [32]. Therefore, we decided to take advantage of the wide variety of publicly available transcriptomic data to explore the relationship between the transcriptomic profiles of genes within the negative and positive pole.

First, we investigated whether the selected genes with similar codon usage could be co-regulated at the transcript level. We analyzed whether changes in abundance of a pair of transcripts from the same pole were correlated with different phenotypic changes. We used transcriptomic data from GTEx and correlated pairwise gene expression across tissues. We compared genes within the positive and the negative poles and between the two poles. The average of pairwise gene expression correlation was significantly higher (Wilcoxon signed-rank test, p≈0) for genes in the negative pole (mean=0.52) than for those in the positive pole (mean=0.18) (Figure 3A). This indicated that the expression of genes in the negative pole (enriched in A/T-ending codons) was coordinated across tissues, but it was not for genes in the positive pole (enriched in G/C-ending codons).

**Figure 3:**
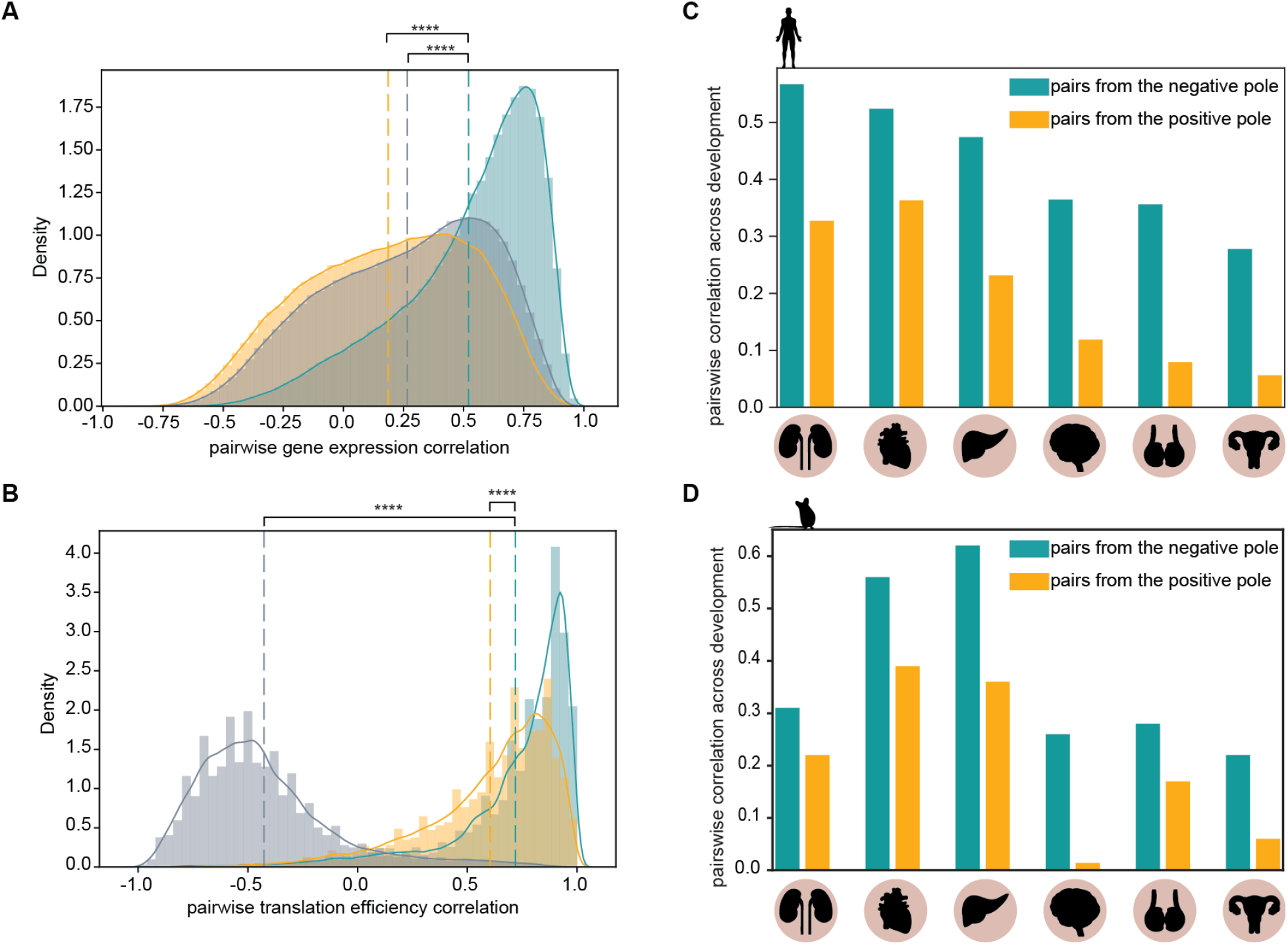
A. Pairwise gene expression correlation across tissues. Genome-wide transcript abundance levels were obtained from GTEX. Distribution of gene expression correlation (Spearman’s ρ) for gene pairs from the negative pole (in blue) were significantly higher (Wilcoxon signed-rank test, p≈0 within numerical error) than correlation for gene pairs from the positive pole (in yellow), and than correlation of pairs of genes with one gene in each of the two poles (in gray) (p≈0). B. Pairwise translation efficiency correlation across tissues. Translation efficiency values were obtained from [33]. Distribution of translation efficiency correlation (Spearman’s *ρ*) for gene pairs from the negative pole (in blue) were significantly higher (Wilcoxon signed-rank test, p≈0 within numerical error) than correlation for gene pairs from the positive pole (in yellow), and than correlation of pairs of genes with one gene in each of the two poles (in gray) (p≈0). C. Average pairwise correlations (Spearman’s ρ) during human development of transcripts of the negative and positive pole. D. Average pairwise correlations (Spearman’s ρ) during mouse development of transcripts of the negative and positive pole.

Then, to explore the effect on translation, we used public ribosome profiling and transcriptomic data from three different human tissues [33], which, in combination, provided a proxy for translation efficiency by the ratio of ribosome footprints to mRNA fragments [34]. We measured pairwise translation efficiency correlations across tissues and observed that both groups of genes tended to have high pairwise translation efficiency correlation (mean > 0.5; Figure 3B). However, genes from the negative pole were significantly (Wilcoxon signed-rank test, p≈0) more coordinated (mean = 0.72) in their translation efficiency than those in the positive pole (mean = 0.60). We also found a significant anticorrelation (mean = −0.42) when the calculation was done among pairs of genes from the two opposite poles, showing an opposite translation profile between poles (Figure 3B).

In RNA-seq data, the technical noise in poorly expressed genes leads to a decrease in the observed correlation compared to highly expressed genes, which are more likely to show a high correlation [35]. Therefore, we controlled for the expression correlation in different bins of gene expression and found that our observations were consistent regardless of the different gene expression levels (Supplementary Figure 2A and 2B). In addition, we performed another control related to the possible coordination of transcripts from genes on the same chromosomes [36]. We found that the correlation between gene pairs of the negative pole was independent of whether they are or not on the same chromosome (Supplementary Figure 2C).

During embryonic development, tightly choreographed abundance changes of hundreds of transcripts are observed. The timely turnover of these patterns of transcripts and corresponding protein products ensure the reliable determination of animal body plans. Therefore, we hypothesized that codon usage is grouping sets of transcripts for coordinated translation, so that genes with similar codon usage will be correlated in their abundances across developmental stages. Taking advantage of the recent expression data on different stages of human development, we explored whether gene coordination was different in the negative and positive poles. We used organ development transcriptomes from humans [37] and found that genes in the negative pole were more coordinated during developmental stages (Figure 3C). This observation was consistent for all six studied organs, although each organ showed a different average correlation, with the kidney presenting the highest correlation. Furthermore, the coordination of this set of protein-coding transcripts was conserved in mouse development (Figure 3D).

Our data demonstrated that the codon usage of a specific gene set enriched in A/T-ending codons played a role in the coordination of their expression in different contexts, for instance, during development.

### Codon usage groups transcripts for developmentally dynamic regulatory processes

Regarding RAS genes and their different codon usage, we investigated whether this opposite codon pattern could lead to opposite changes in expression during developmental stages. Interestingly, we observed that the transcript abundance of KRAS and HRAS alternates during development (Figure 4A). A compensatory switch in the expression of these paralogous genes could indicate a role in keeping a constant essential function for cell viability while having differential functionalities, which serve a specific role. Then, we investigated other families with members having an opposite codon usage (highest and lowest quartile of the PC1: >3.25; <-5. 14) and a different codon conservation in mammals (higher and lower quartile of the conservation: >0.85; <0.80) (Figure 4B), as observed for KRAS and HRAS. We found seven other pairs of paralogs, with CALM2 and CALM3 as a paradigmatic example: both genes coded for the exactly same protein (100% amino acid identity) but had a completely different codon usage, since CALM2 but not CALM3 belonged to the negative pole in the PC1 analysis. On average, we found the same pattern of switching during development when considering all the families together (Figure 4C). The standard deviation of the family members in the positive pole was more important than for the negative, highlighting the lower coordination globally observed for genes of the positive pole.

**Figure 4:**
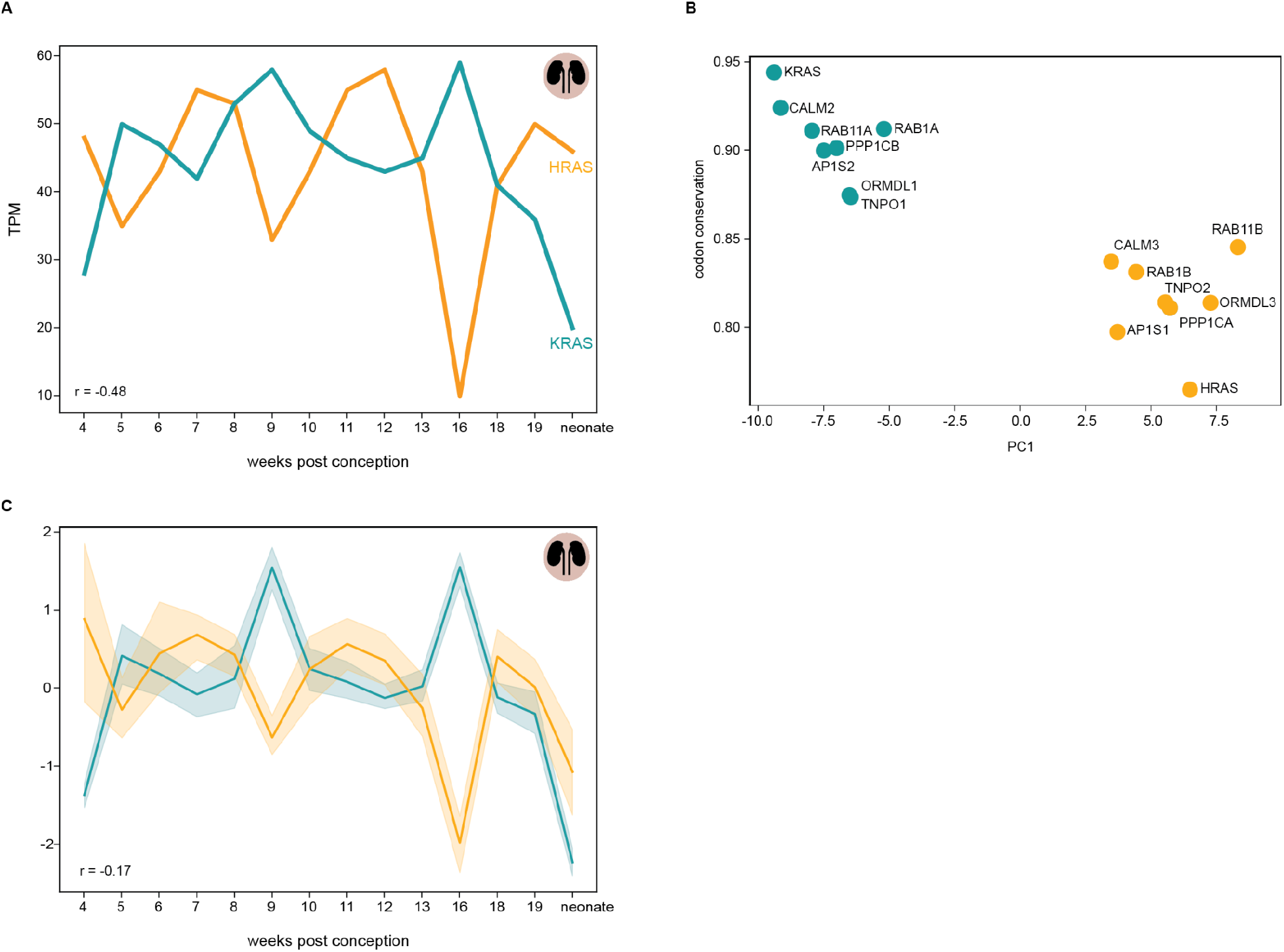
A. KRAS and HRAS transcripts turnover during kidney development. B. KRAS and HRAS are found at opposite ends of codon usage (PC1) and codon conservation axes. We found seven other families with the same pattern. C. Relative average changes in transcript abundance during kidney development for the eight family members of the negative and the positive pole.

### Similar patterns of codon usage lead to interacting proteins

The above-mentioned data suggested a coordinated regulation of the expression and translation of genes in the negative pole, and a weaker correlation for those in the positive pole. Hence, we evaluated whether genes need to be expressed coordinately in order to function in the same time and space, as for example to function in the same protein complexes [38]. To that end, we calculated the PC1 distances of genes belonging to the same protein complex [39] and compared them with the distances of genes not belonging to the same complex. We found that genes encoding proteins that are part of the same protein complex had significantly smaller PC1 distances, meaning similar patterns of codon usage (Wilcoxon signed-rank test, p=1.2e-72) (Figure 5A). To the best of our knowledge, this was the first description of this phenomenon in human proteins, and it has been very recently observed in bacteria [40, 41]. Then, we analyzed whether genes found in the negative pole (more coordinated in translational efficiency) showed a higher tendency to form complexes. Indeed, proteins coded by genes in the negative pole were consistently found in more complexes than genes in the positive pole (Figure 5B). These results indicated that their codon usage helped coordinate their abundance, which in turn allows a coordinated formation of functional protein complexes.

**Figure 5:**
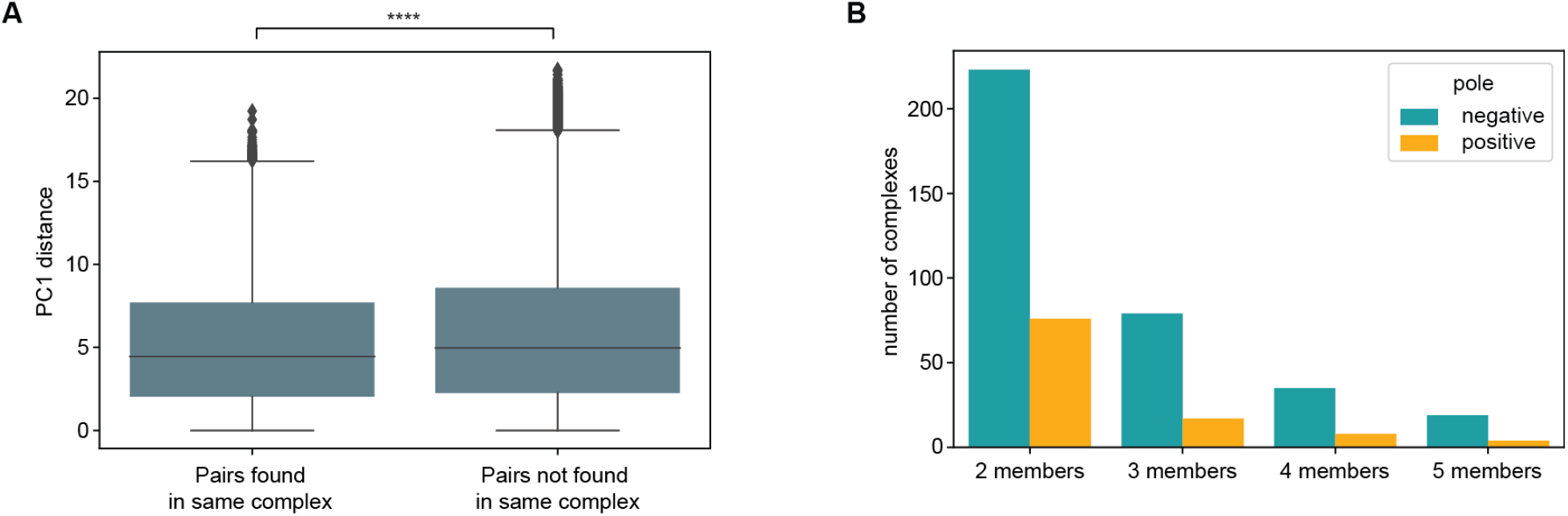
A. PC1 distances of gene pairs of the same or different protein complex (Wilcoxon signed-rank test, p=1.2e-72). B. Comparison of the amount of complexes formed by the protein in the negative and the positive pole.

### tRNA expression profiles

The main feature of the codon usage of genes found in the negative pole was their enrichment in A/T-ending codons. Thus, we hypothesized that tRNAs with ANN or UNN anticodons could contribute to the coordinated changes in transcript abundance of the genes in the negative pole. The observed coordination could reflect a greater variation of the tRNA pool related to A/T-ending codons than that of G/C-ending codons. Harnessing recently published data on tRNA abundance [22], we explored the variation of tRNA across healthy tissues. We found that the coefficient of variation among tissues was higher for the pool of tRNAs with A or U in the position 34 of the anticodon than for tRNAs with G or C (Figure 6A, Supplementary Table 8). Our analysis showed that the fine-tuned regulation of transcripts enriched in A/T-ending codons could be directly related to the specific regulation of tRNAs with anticodons recognizing A/T-ending codons. Therefore, we used the Supply-to-Demand Adaptation (SDA) codon values to obtain a global estimate of translation efficiency [22]. We compared the SDA values from healthy samples for codons enriched in the negative pole component against those in the positive pole (see Figure 2D). We observed that codons enriched in the negative PC1 (A/T-ending codons plus TTG and AGG) correlated more among tissues than the ones in the positive pole (G/C-ending codons except TTG and AGG) (Wilcoxon signed-rank test, p=3.5e-76) (Figure 6B). This observation suggested that A/T-ending codons were more coordinated in their translation efficiency than G/C-ending codons in relation to tRNA abundances. Interestingly, there was a significant anticorrelation between the two groups of codons (Wilcoxon signed-rank test, p=9.7e-73).

**Figure 6:**
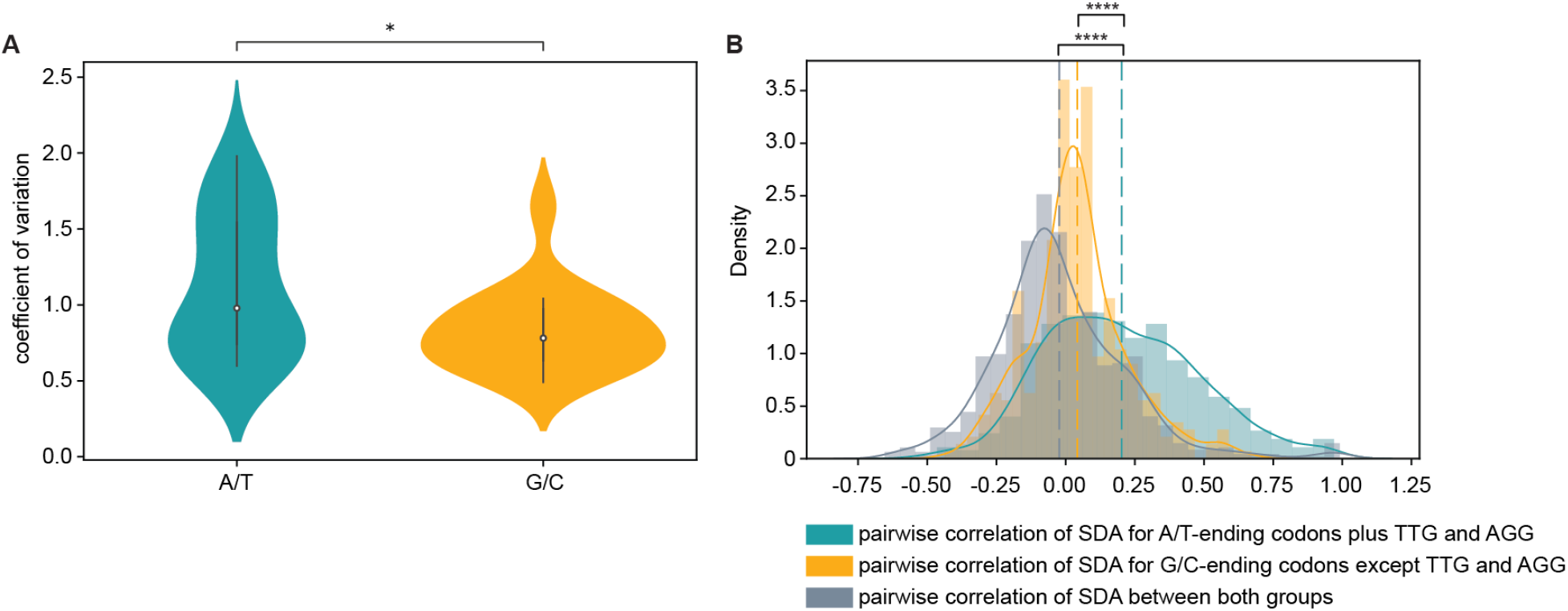
A. Average of coefficient of variations for the expression of tRNAs with anticodons recognizing A/T-ending codons (blue) and G/C-ending codons (yellow) (t test, *P < 3.704e-02). B. Pairwise correlations (Spearman’s ρ) of SDA values for two groups of codons “A/T-ending codons plus TTG and AGG” and “G/C-ending codons except TTG and AGG”.

### Codon differences in mammals

To gain further insight into codon conservation, we investigated the type of codon differences observed between different mammals. Since A and T-ending codons were enriched in the genes with high codon conservation across mammals, we hypothesized that this conservation is connected to the need of these genes for co-regulation, probably with a finely tuned mechanism related to changes in tRNA abundance. Therefore, we expected that the main codon differences observed between humans and mammals would involve keeping A- or T-ending codons in genes of the negative pole. By counting the average synonymous differences for A-, T-, C- or G-ending codons (e.g., number of differences for GGC in chimpanzee compared to GGA in human), we found that the coding sequences in the negative pole tended to have more differences for A and T than G and C (Supplementary Figure 3A and B). Specifically, a T in a particular species was seen more frequently as an A, than as a G or C in other species, while a G was seen more frequently as an A, or a T, than as a C. The opposite trend was observed for genes in the positive pole. The same pattern was unveiled when we compared to other distant mammals, such as chimpanzee, dog, and armadillo (Supplementary Figure 3A and B).

### Codon usage variations in vertebrates and non-vertebrates

Subsequently, we explored the codon usage trends in different species of vertebrates and non-vertebrates. We first applied principal component analysis (PCA) to the relative codon usage frequencies of all individual genes for eight different vertebrates (Supplementary Figure 4) and seven non-vertebrates (Supplementary Figure 5). We observed that the distribution of codons across PC1 components was similar in all species, especially the first axis of the PCA separating A/T- and G/C-ending codons was consistent across species (Figure 7A; Supplementary Table 9).

Since the polarization between genes with A/T- or G/C-ending was deeply rooted in vertebrate evolution, we investigated how our two groups of studied human genes in the negative and positive poles projected across the first two PCs of the different vertebrate and non-vertebrates species by using their orthologs (Figure 7B and Supplementary Figures 4 and 5). As above-mentioned, human genes in the negative pole are enriched in A/T-ending codons. This enrichment was found to be similar in all analyzed vertebrate species, except for *Danio rerio* (Figure 7B, Supplementary Figure 4). This could be related to the fact that teleost fish underwent a third round of whole genome duplication compared to the two rounds of vertebrates [42]. This enrichment was lost when looking at non-vertebrates (Figure 7B, Supplementary Figure 5). Interestingly, the distribution of genes in the positive pole widens across species, highlighting the lower codon conservation of these genes (Figure 7B). Additionally, the explained variance of the PC1 decreases (Supplementary Figures 4 and 5), showing a reduced codon bias in the genomes of invertebrates compared to those of vertebrates. We observed comparable patterns with one-to-one pairwise orthologs (both genes in the pair had only one ortholog in the other species) (Supplementary Figures 6 and 7).

**Figure 7:**
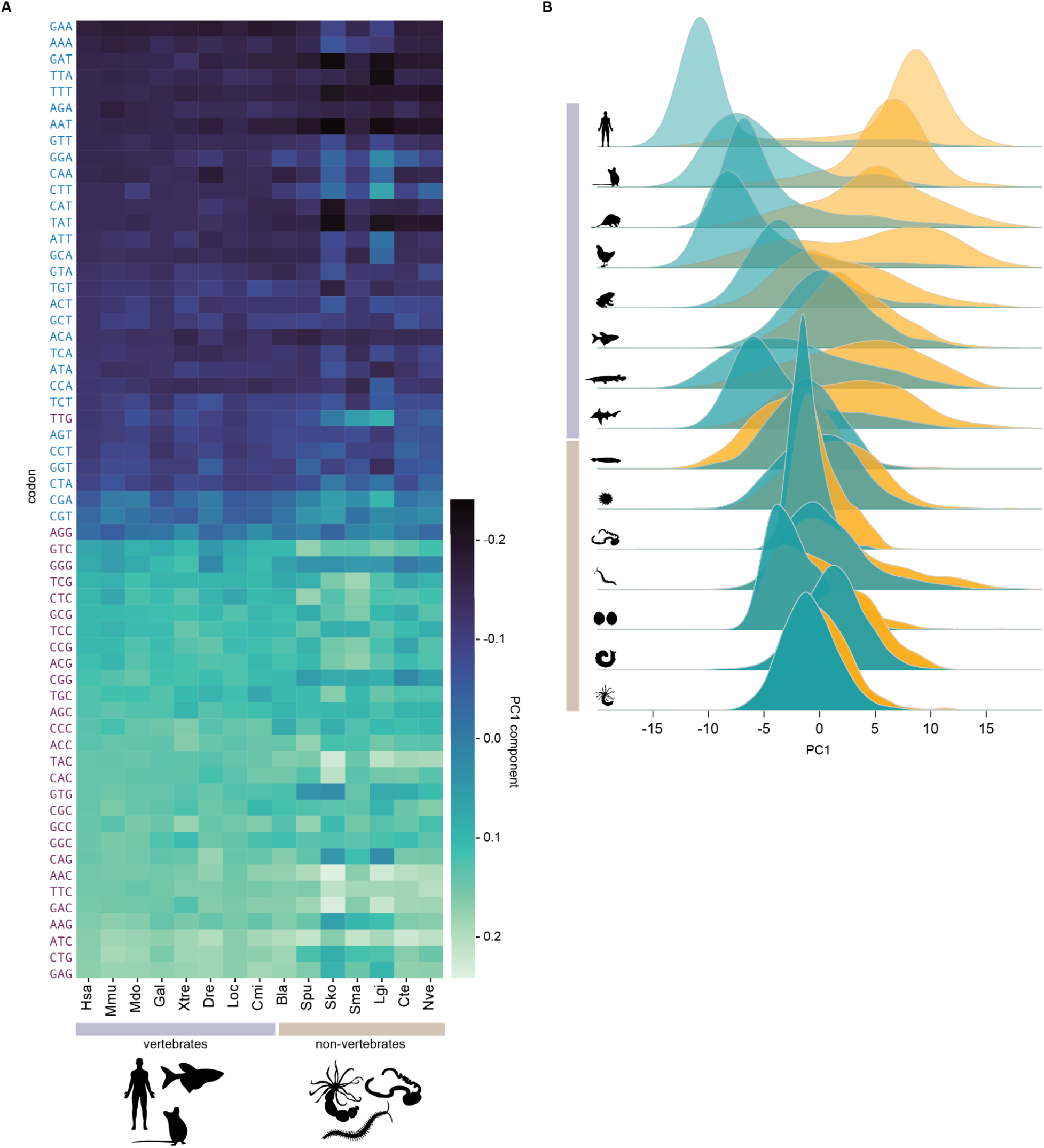
We analyzed eight vertebrate and seven invertebrate species. (Hsa: Homo sapiens, Mmu: Mus musculus, Mdo: Monodelphis domestica, Gal: Gallus gallus, Xtre: Xenopus tropicalis, Dre: Danio rerio, Loc: Lepisosteus oculatus, Cmi: Callorhinchus milli, Bla: Branchiostoma lanceolatum, Spu: Strongylocentrotus purpuratus, Sko: Saccoglossus kowalevskii, Sma: Strigamia maritima, Lgi: Lottia gigantea, Cte: Capitella teleta, Nve: Nematostella vectensis). A. Codons are ordered according to their coefficient of the first eigenvector (PC1) in humans. The heatmap highlights the changes of this order in other species. Note that the scale of the values is arbitrary, as only the relative values are important (direction of the vector in the multidimensional space). Negative values indicate a negative PC1 and vice versa. B. Distribution of the ortholog genes in the negative (blue) and the positive pole (yellow) along the main codon usage axis revealed a polarization for most of the species of vertebrates but not for non-vertebrates. Explained variances can be found in Supplementary Figures 5 and 6.

These data highlighted the separation between genes with A/T- or G/C-ending in other vertebrates, suggesting that the expression of these orthologs might also be coordinated in species other than mammals.

## Discussion

Since the discovery of the genetic code, specific protein sequences are known to be potentially encoded by various synonymous alternatives. These synonymous variants have been proposed to be the result of the fine interplay between mutational biases and selection processes. Aside from their origin, their functional implications have been also investigated; whether and how codon usage preferences can be linked to specific regulatory processes such as modifying mRNA stability [12] or protein folding [16]. In the present work, we showed the connection between codon usage and the coordination of mRNA pools in different cellular contexts.

Here, we observed a general biased codon usage for human genes with synonymous codon conservation. For instance, our results showed that the three RAS paralogous genes, which display differential codon usage, had a distinct codon conservation among mammals. This was also the case for other gene families, such as the striking case of CALM2 and CALM3, which code for a protein 100% identical but had different codon bias and codon conservation. Specifically, protein-coding genes with a conserved codon usage (e.g., KRAS and CALM2) were enriched in A/T-ending codons.

As previously shown, genes with similar codon usages should have similar expression patterns, albeit displaying different average expression levels [43]. This suggests that codon usage can serve as a molecular switch to alter expression programmes to meet the cell’s requirement. The adaptation between codon choice on mRNAs and tRNAs pools has a dual regulatory role in expression, as it appears to determine not only translation efficiency but also transcript stability [12, 32]. Our analysis revealed that protein-coding transcripts enriched in A/T-ending codons were coordinated in their expression patterns across tissues or within different stages of mammalian ontogenesis. Additionally, we found that proteins forming complexes were generally encoded by genes enriched in A/T-ending codons, with codon bias possibly impacting the topology of the protein-protein interaction networks. The role of this coordination could be related to the association of transcriptomes enriched in A/T-ending codons to specific cellular contexts, such as rapidly dividing cells [20, 44, 45] or under splicing regulation [13]. However, this coordination has not been observed in transcripts enriched in G/C-ending codons, such as housekeeping genes that are expected to maintain constant expression levels in all conditions [46] or genes involved in differentiation [20]. We speculated that this coordination could be achieved through the orchestration of tRNA expression and tRNA modification [47]. Indeed, we found that the expression variance across healthy tissues of tRNAs with A or U in position 34 of the anticodon was higher than that for G or C. In addition, based on tRNA abundances, we found that A/T-ending codons had a coordinated translation efficiency. Therefore, our observations would point towards a mechanism that fine-tunes the coordination of A/T-ending codon enriched transcripts and that could be conserved in mammals. Also, we interpret this as the necessity for A/T-ending codons to be conserved, since they are linked to specialized spatio-temporal expression (e.g., tissue-specificity or stage-related transcript turnover). Indeed, proteins with highly specific roles have been shown to be modulated more frequently than those with central functions [48]. On the other hand, G/C-ending codons would be subjected to a lower selection, as they would be related to a robust and less variable tRNA expression that would facilitate broad expression across all types of cellular context. However, the antagonistic expression pattern of paralogous genes, for example, during development, suggest that other possible mechanisms should be disentangled. Since during development some tissues present larger differences between transcript pairs than others, it would be interesting to know how this correlates with the magnitude of tRNAs variation in the different tissues across development. The observed codon conservation in genes of the negative pole of the PCA led us to speculate that mammal genomes have highly similar tRNA regulation to decode this group of genes. Particularly, if tRNA profiles were changing considerably, different codons would be selected in each organism. Even though in recent years evidence has pointed towards the interplay between codons and tRNA availability as a means to dynamically regulate protein abundance, the mechanisms behind this regulation remain to be elucidated. Several hypotheses suggest that individual tRNAs are specifically modulated in response to various environmental and physiological cues to support specific roles in a large number of biological processes. How tRNA-specific transcription is established is starting to be explored in mammalian cells, for instance by studying signals of chromatin marks associated with activation [20], Pol II and Pol III tRNA-specific transcription events [49], or SOX4-dependent regulation of a subset of tRNA genes [50].

Intriguingly, the codons TTG-Leu and AGG-Arg were G-ending codons systematically clustering with A/T-ending codons in our analyses (Figure 2D, Figure 7A). The reasons underlying this observation were unclear. Indeed, arginine and leucine allow GC-changing synonymous substitutions in the first and third codon positions and have codons which may be expected to show different usage patterns.

Our results included other organisms, especially vertebrates. We observed that the pattern between A/T- and G/C-ending codons was preserved across divergent species of vertebrates and to a lesser extent in non-vertebrates. Also, orthologs of genes of the negative pole were clustering in other species of vertebrates, suggesting that the mechanism regulating the coordination of these genes would not only be present in mammals but also in other species of vertebrates.

## Conclusions

Taken together, our work underlies codon conservation due to selection for expression coordination. Further investigation will be needed to better understand the significance and implications of the findings presented here. We are now entering an era in which tRNA abundances and tRNA modifications can be quantified with precision in many cell types and organisms and across different timescales and conditions. Expanding further our knowledge regarding codon signatures and the dynamics of the tRNA pool will have a profound impact on our understanding of gene expression regulation.

## Methods

### Coding sequences

The coding sequences of *Homo sapiens* were downloaded from the Consensus CDS (CCDS) project (ftp://ftp.ncbi.nlm.nih.gov/pub/CCDS/), release 2016/09/08.

### Multiple sequence alignment

The alignments were downloaded using CodAlignView [51] by specifying the coordinates of all human coding sequences obtained from the CCDS.

### Amino acid conservation

We used the multiple sequence alignments obtained with CodAlignView, which aligns the human sequence to the corresponding sequences of mammals. Then, for each species, we compared each amino acid of the sequence to that of human. We calculated the identity taking into account the amount of amino acids conserved between human and the mammal, and we normalized the total amount of considered amino acids (conserved amino acids + non-conserved amino acids). Then, we averaged the identity value of all mammals (Supplementary Table 1).

### Codon conservation

Similar to amino acid conservation, we measured codon conservation using the multiple sequence alignment from CodAlignView that aligns the human sequence to the corresponding sequences of mammals. Then, for each species, we compared each codon of the sequence to that of human. We calculated the identity taking into account the amount of codons conserved between human and the mammal and we normalized by the total amount of considered codons (conserved codons + codons with synonymous changes) (Supplementary Table 2). Non-synonymous changes were ignored. Then we averaged the identity value of all mammals.

### Codon usage PCA and conservation

For the following analysis, we selected only the CCDS that had an amino acid conservation among mammals higher than 0.90. Also, we kept one transcript variant per gene. We applied PCA to the relative synonymous codon frequencies [52] of all individual human coding sequences. We defined our PCA projection based on the codon usage distribution of individual genes (as previously reported [21]). When computing the PCA of individual genes, we first excluded single codon families (AUG and UGG). Second, for coding sequences that lacked codons of a specific family, we imputed values with the average codon frequency across all genes. Finally, we colored each gene according to its codon conservation (Supplementary Table 3).

### GO enrichment analysis

We downloaded the human gene ontology annotation from Ensembl (v102). We performed a gene ontology enrichment analysis on the genes of the negative and positive poles (Supplementary Table 5) with the R package TopGO, using as a background all genes with a functional annotation. P values correspond to uncorrected two-sided Fisher’s exact tests as provided by TopGO (Supplementary Table 6).

### GC3 content

GC3 content was calculated for all studied coding sequences (Supplementary Table 4).

### Pairwise correlations

Pairwise gene expression correlation (Spearman’s *ρ*) coefficient values were measured from the transcriptomic and ribosome profiling data. Correlations were obtained across tissues for GTEx and translation efficiency data for three different tissues [33] and across developmental stages [37].

### Protein complexes

The known human protein complexes were obtained from the CORUM database [39] (Corum 3.0 current release, September 2018). For each protein pair found together in at least one protein complex, we calculated the PC1 distance (PC1_gene1_-PC1_gene2_=PC1 distance) of the corresponding genes and compared them with protein pairs not found in the same complex. Also, we compared the number of complexes with two to five proteins belonging to the same complex of the negative or positive pole.

### Paralogs Ensembl

To define gene families, we retrieved information regarding protein sequence similarity and family membership from Ensembl. Ensembl’s family classification often contains outliers that have a much lower sequence similarity than other proteins of the same family. As a larger variation in amino acid sequence implies greater variation of biochemical function, and because in our study we aimed to focus on differences in codon usage, we sought to remove these outliers. We therefore applied another, more stringent filter; for each family, we computed the similarity distribution of all members to a consensus member. We then removed all family members that had a lower similarity than the mean similarity minus one standard deviation.

### tRNA abundances

tRNA abundances and Supply-to-Demand (SDA) values from healthy samples were obtained from previously published work [22].

### Codon differences among mammals

We measured codon conservation using the multiple sequence alignment from CodAlignView that aligns the human sequence to the corresponding sequences of mammals. Then, for each species, we compared each codon of the sequence to that of human, chimpanzee, dog, or armadillo, and counted the number of synonymous changes between codons differing only in the third nucleotide position (i.e., the wobble position). To normalize changes within codon boxes, we computed the percentage of 3^rd^-nt changes in human, chimpanzee, dog, or armadillo with regard to the total of changes of each codon in each species, considering only codon boxes with a total of at least 3 changes. Finally, we averaged the relative codon changes across all mammals.

### Orthology analysis

We ran Broccoli v1.2 [53] to infer gene orthogroups among all protein-coding genes in a set of eight vertebrate and seven invertebrate species, only considering one representative protein isoform per gene (i.e., the one with the longest coding sequence). We used the gene annotations for the following assemblies and versions from Ensembl (https://www.ensembl.org/): human *(Homo sapiens:* hg38, v88), mouse *(Mus musculus:* mm10, v88), opossum *(Monodelphis domestica:* monDom5, v86), chicken *(Gallus gallus:* galGal6, v99), clawed frog *(Xenopus tropicalis:* XenTro9, v101), zebrafish *(Danio rerio:* danRer10, v80), spotted gar *(Lottia gigantea:* LepOcu1, v88), and elephant shark *(Callorhinchus milii:* 6.1.3, v99). We used the gene annotations for the following assemblies and versions from Ensembl Metazoa (https://metazoa.ensembl.org/): centipede *(Strigamia maritima:* Smar1, v26), owl limpet *(Lottia gigantea*: Lotgi1, v40), polychaete annelid worm (*Capitella teleta*: capTel1, v45), sea anemone *(Nematostella vectensis:* NemVec1, v36). Plus, we considered amphioxus *(Branchiostoma lanceolatum:* braLan3, from ucsc.crg.eu), sea urchin *(Strongylocentrotus purpuratus:* strPur4, from SpDB), and acorn worm *(Saccoglossus kowalevskii:* sacKow3, from BCM-HGSC). All orthologs of human genes of the positive or negative pole in each species were considered in the analysis.

## Declarations

### Ethics approval and consent to participate

Not applicable

### Consent for publication

Not applicable

### Availability of data and materials

The datasets analyzed during the current study are available in the Supplementary Table 10. The Genotype-Tissue Expression (GTEx) Project was supported by the Common Fund of the Office of the Director of the National Institutes of Health, and by NCI, NHGRI, NHLBI, NIDA, NIMH, and NINDS. The data used for the analyses described in this manuscript were obtained from the GTEx Portal accession number phs000424.v8.p2. All data generated during this study are included in this published article and its supplementary information files. The scripts are reported in the Github repository: https://github.com/HannahBenisty/CodonUsage.git

### Author Contributions

Conceptualization: H.B; Methodology: H.B, X.H-A, M.W, M.A-G, F.M, L.R, G.S, D.W, M.I, M.S, L.S; Validation: H.B, X.H-A, M.W, M.A-G, F.M, L.R, G.S, D.W, M.I, M.S, L.S; Formal analysis: H.B, X.H-A, M.W, M.A-G, F.M, L.R, G.S, D.W, M.I, M.S, L.S; Writing-Original Draft: H.B; Writing-Review & Editing: H.B, X.H-A, M.W, M.A-G, F.M, L.R, D.W, M.I, M.S, L.S; Visualization: H.B; Funding Acquisition: L.S.; Supervision: L.S.

## Acknowledgments

We thank Juan Valcárcel and Fran Supek for their comments and suggestions. We acknowledge the support of the Spanish Ministry of Economy, Industry and Competitiveness (MEIC) to the EMBL partnership, the Centro de Excelencia Severo Ochoa and the CERCA Programme / Generalitat de Catalunya.

---

**Supplementary Figure 1:**
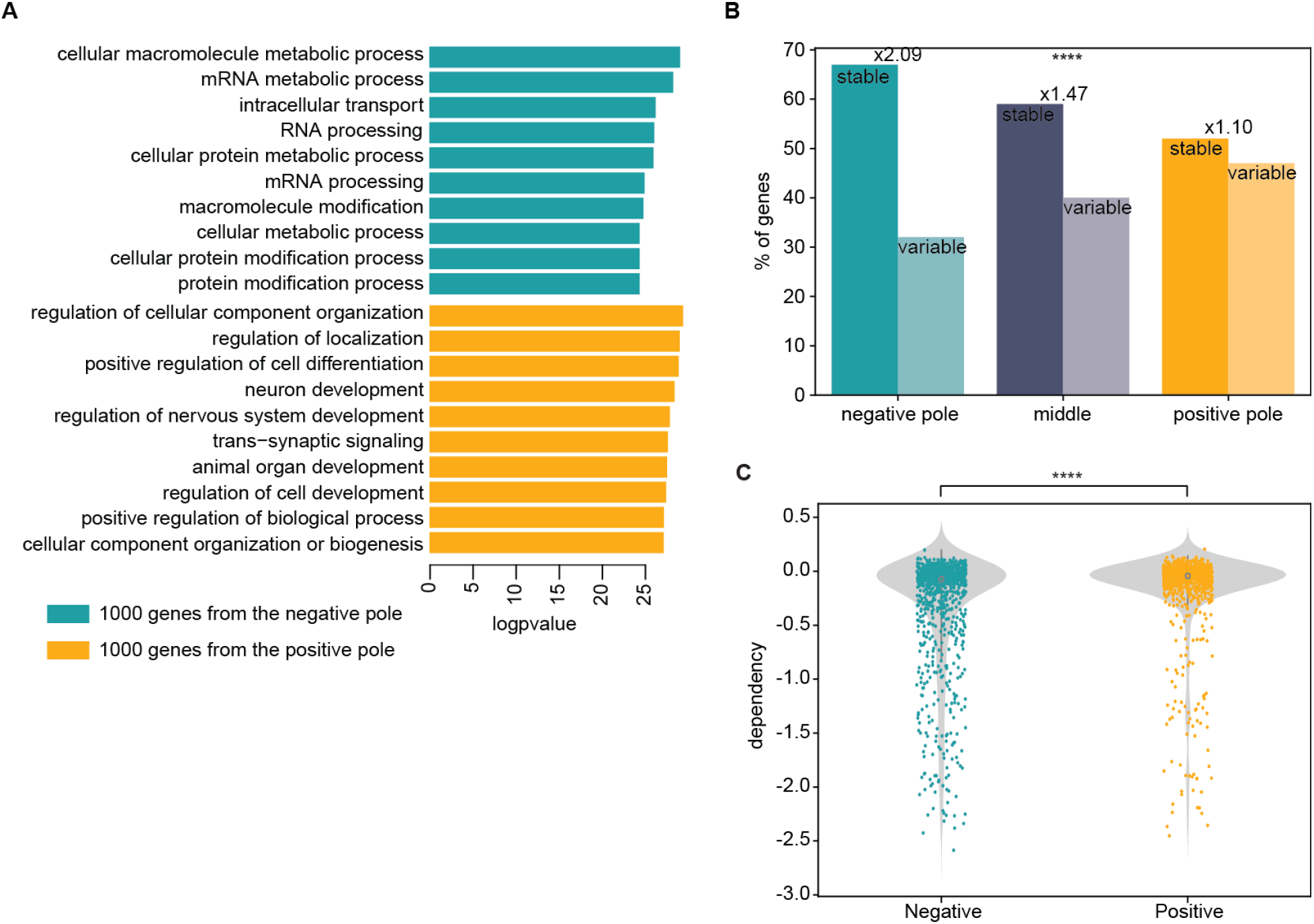
A. Comparison of gene ontology enrichment analysis of genes in the two opposite poles. GO analysis is conducted for each group through biological process enrichment. Only the 10 most significantly enriched GO terms (FDR < 0.05) are shown. B. Percentage of genes within poles that correspond to genes with expression levels that are highly (stable) or poorly (variable) correlated among mammalian species [27] ****P < 0.0001, Chi-squared test. C. Pan-cancer essentiality scores in DepMap database from the genes in the negative and positive poles (Wilcoxon signed-rank test, p= 2.9e-10).

**Supplementary Figure 2:**
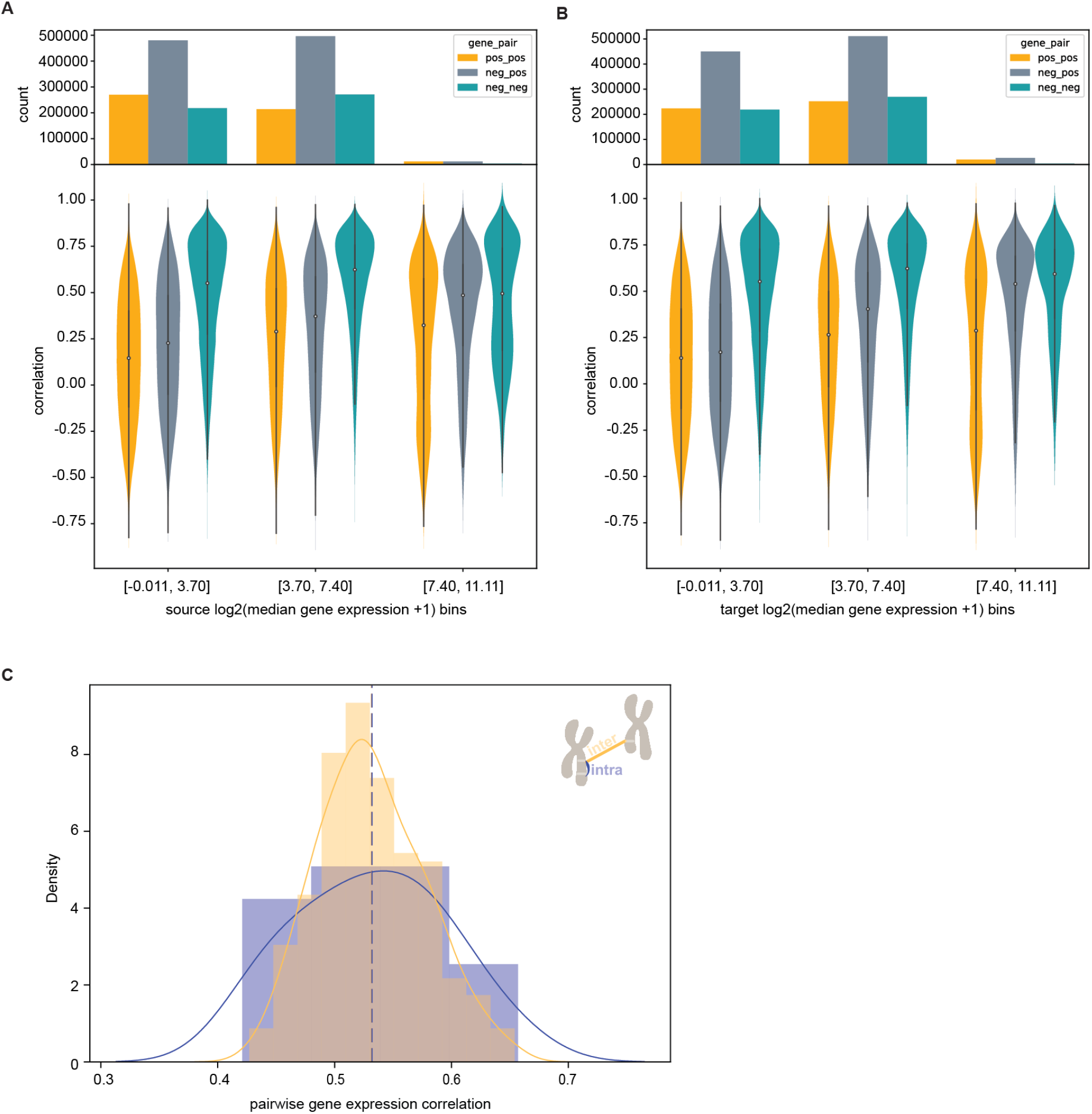
A-B. Pairwise gene expression correlation (Spearman’s *ρ*). Genome-wide transcript abundance levels were obtained from GTEX. Gene pairs from each pole are correlated (gene pairs of the negative pole in blue and those of the positive pole in yellow). Correlation of gene pairs among the two poles in gray. Bins were built based on the gene expression of source genes (A) or target genes (B). C. Pairwise gene expression correlation for genes in the negative pole. Gene correlation for gene pairs belonging to the same chromosome in purple and correlation for genes from different chromosomes in yellow.

**Supplementary Figure 3:**
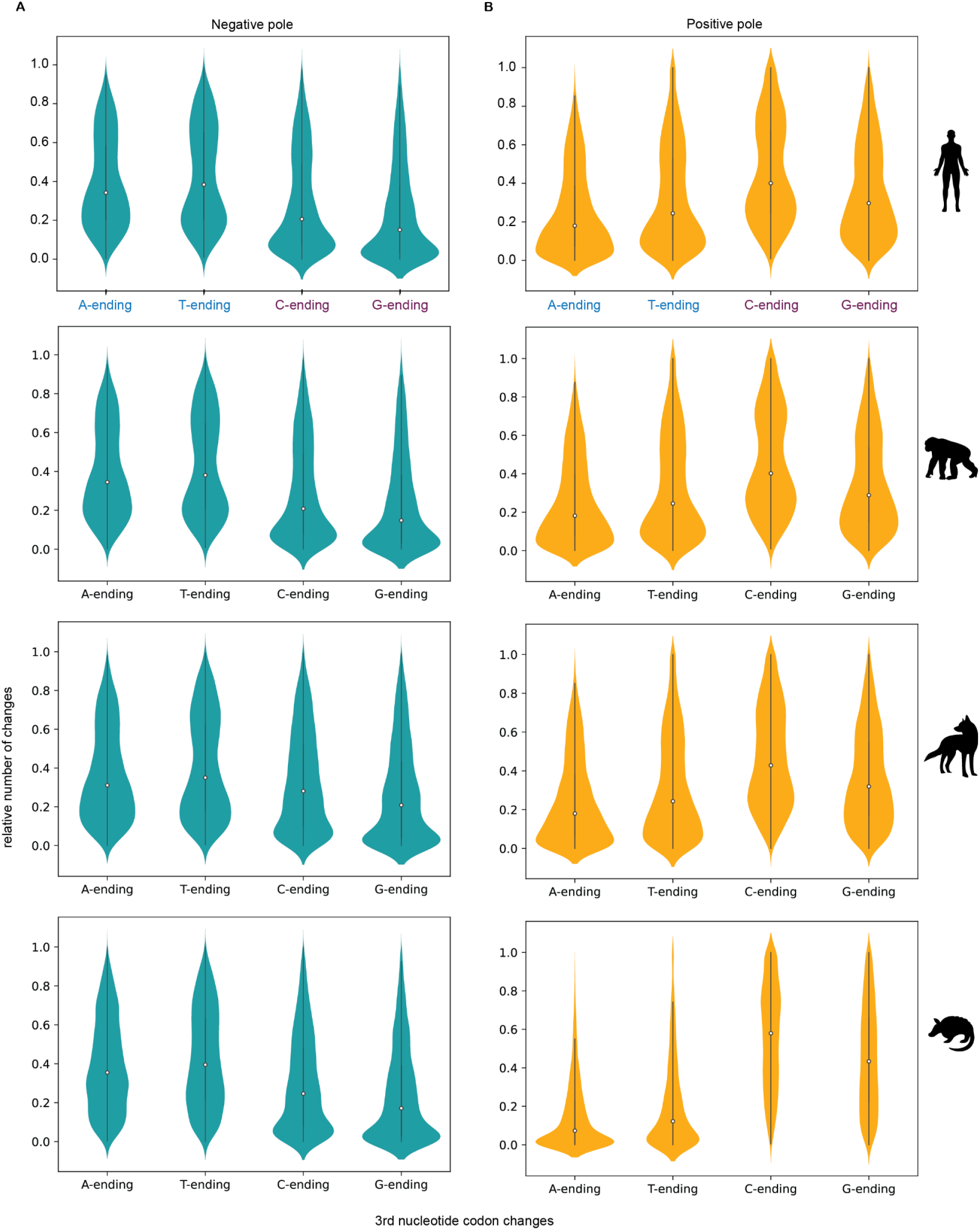
A. Average of 3rd base codon changes for A-, T-, C- or G-ending codons in the coding sequences of the negative pole in human, chimpanzee, dog, and armadillo. B. Average of 3rd base codon changes for A-, T-, C- or G-ending codons in the coding sequences of the positive pole in human, chimpanzee, dog, and armadillo.

**Supplementary Figure 4:**
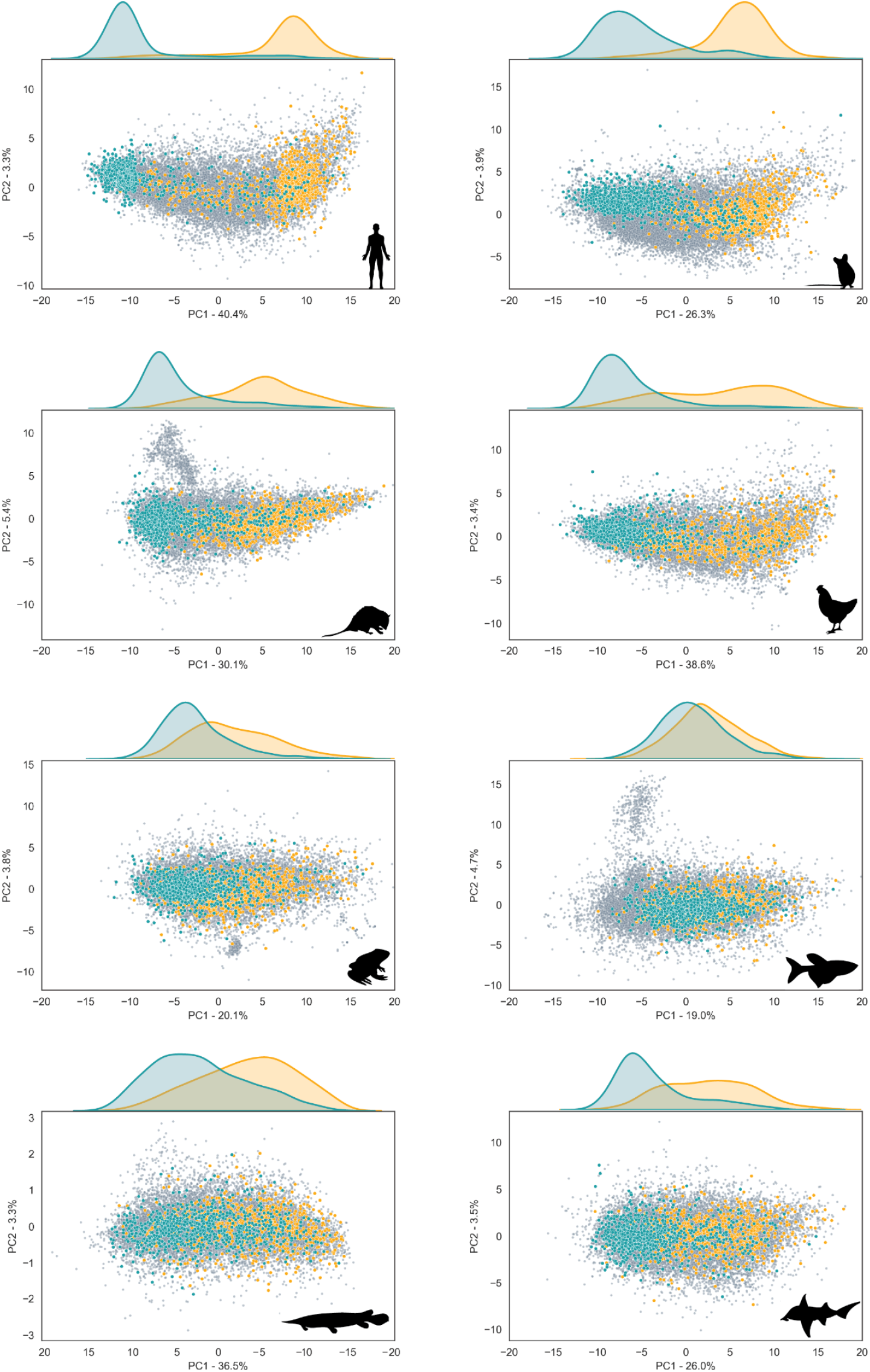
PCA projections of the codon usage of genes for eight different vertebrates. The location of each gene was determined by its codon usage. Distribution along the main codon usage axis revealed a polarization for most species: genes in the negative pole (blue) and those in the positive pole (yellow).

**Supplementary Figure 5:**
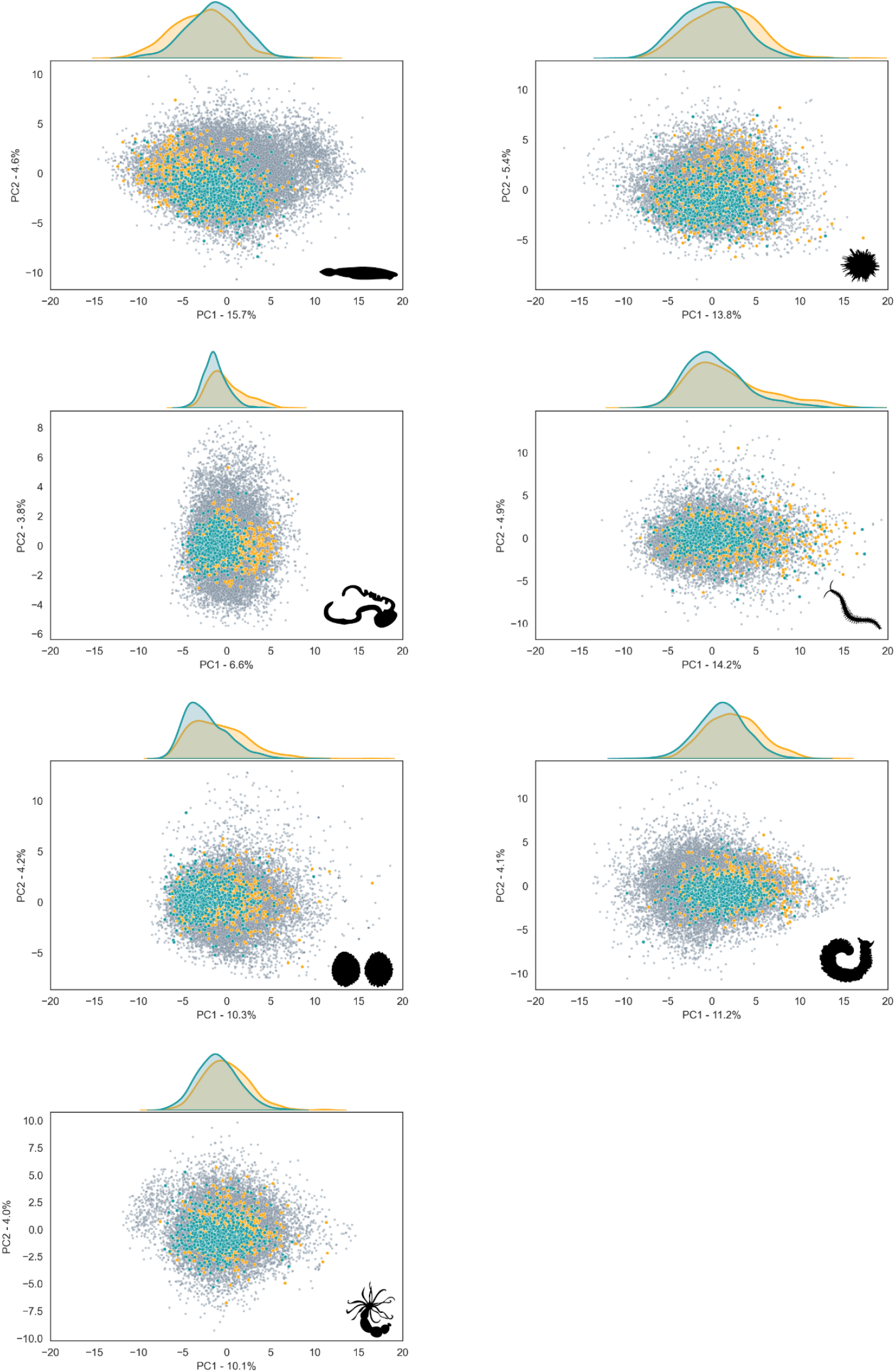
PCA projections of the codon usage of genes for seven different non-vertebrates. The location of each gene was determined by its codon usage. Distribution along the main codon usage axis revealed a polarization for most species: genes in the negative pole (blue) and those in the positive pole (yellow).

**Supplementary Figure 6:**
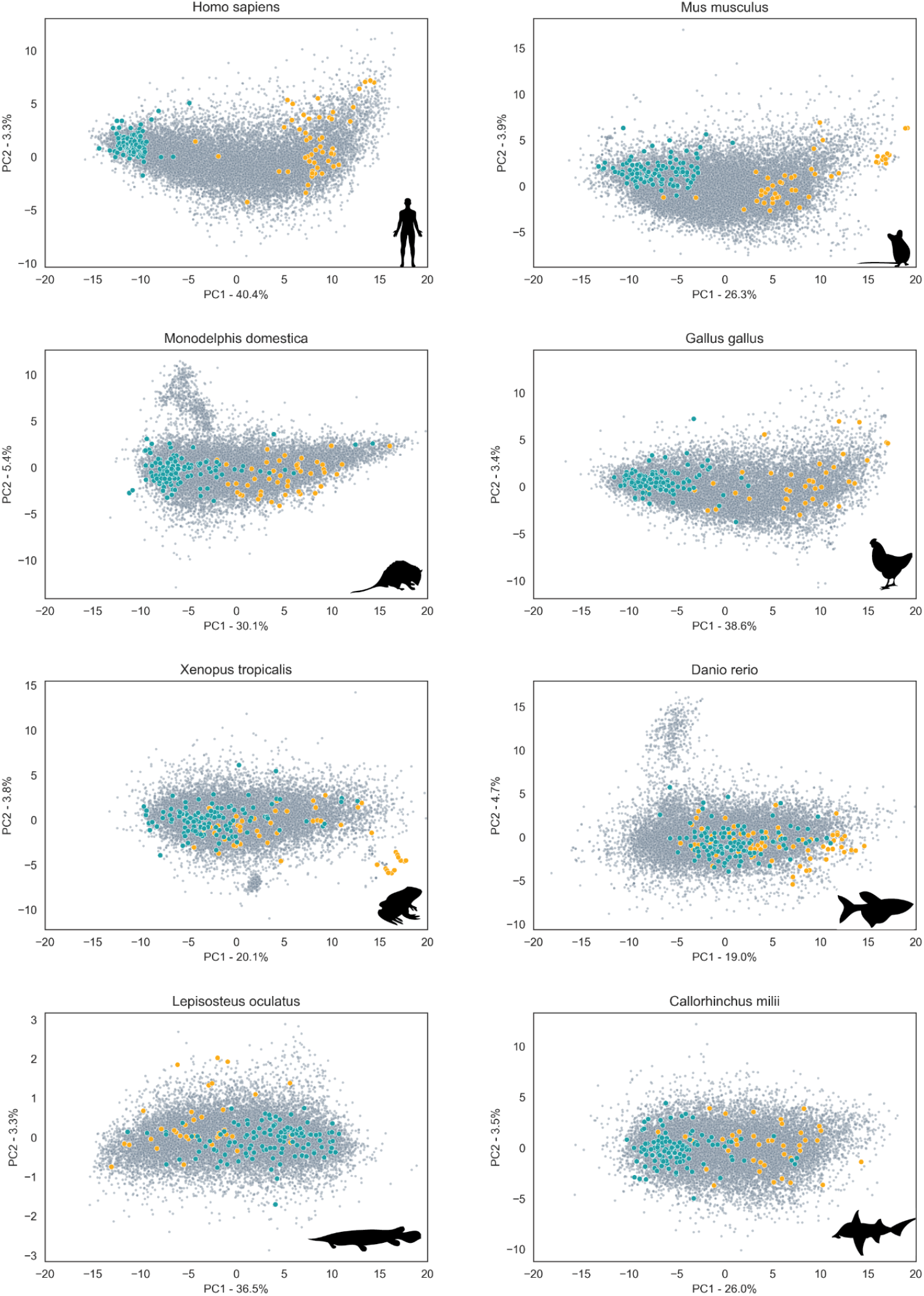
PCA projections of the codon usage of genes for eight different vertebrates for one-to-one pairwise orthologs. The location of each gene was determined by its codon usage. Distribution along the main codon usage axis revealed a polarization for most species: genes in the negative pole (blue) and those in the positive pole (yellow).

**Supplementary Figure 7:**
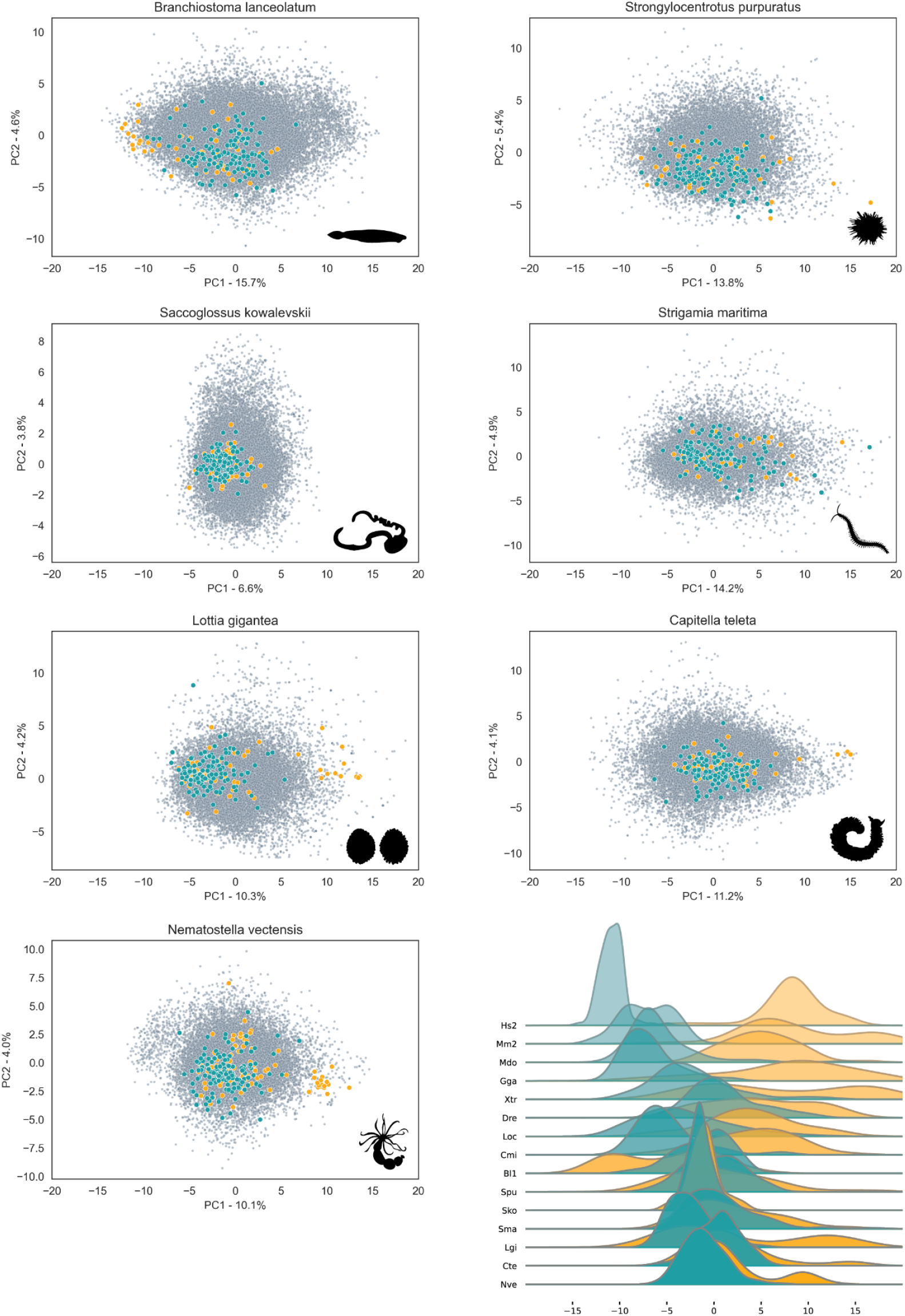
PCA projections of the codon usage of genes for eight different non-vertebrates. The location of each gene was determined by its codon usage. Distribution along the main codon usage axis revealed a polarization for most species: genes in the negative pole (blue) and those in the positive pole (yellow). The last panel corresponds to the distributions of the ortholog genes in the negative pole (blue) and in the positive pole (yellow) along the main codon usage axis. A polarization for most species of vertebrates but not for non-vertebrates was revealed.

